# *In vitro* hepatotoxicity assessment of *Lippia javanica* (Burm.f.) Spreng. aqueous leaf extract

**DOI:** 10.1101/2021.05.26.445727

**Authors:** Bresler Swanepoel, Trevor C. Koekemoer, Luanne Venables, Elsabe Cloete, Nonhlanhla P. Khumalo, Maryna van de Venter

**Author notes:** Corresponding author: Department of Biochemistry and Microbiology, PO Box 77000, Nelson Mandela University, Port Elizabeth, South Africa 6031. Email addresses (Bresler Swanepoel), (Trevor C. Koekemoer), (Luanne Venables), (Elsabe Cloete), (Nonhlanhla P. Khumalo), (Maryna van de Venter).

## Abstract

**Ethnopharmacological relevance:** *Lippia javanica* leaves are popular in traditional food, medicine and for insecticidal uses in various Africa countries and North-East India. Anecdotal evidence suggests that it is safe to use but limited animal studies suggested potential toxicity at high dosages, including hepatotoxicity.

**Aim of the study:** To screen for potential hepatotoxicity of *L. javanica* leaf extracts *in vitro*, thereby contributing to its toxicological profile for safe use in food and topical applications.

**Materials and methods:** High content analysis techniques and fluorescent dyes were used to monitor C3A hepatocarcinoma cells for changes in morphological features that are associated with development of mitotoxicity, steatosis, oxidative stress, and lysosomal dysfunction.

**Results:** No changes were observed in cell viability, reactive oxygen species or lysosomal content at concentrations up to 200 µg/ml in C3A cells. Mitochondrial membrane potential was reduced by approximately 10% but this effect was not dose-dependent nor was it accompanied by a reduction in mitochondrial content. A dose-dependent decrease was observed in neutral lipid content.

**Conclusion:** The results from this *in vitro* study suggest that *L. javanica* leaf extracts is not anticipated to be hepatotoxic at concentrations in the range that is assumed for food or topical use.

## 1. Introduction

*Lippia javanica* (Burm.f.) Spreng. (Verbenaceae) has a long traditional use in human welfare. Leaves are cooked as vegetable in North-East India (Basumatary et al., 2015; Brahma et al., 2013); are used as food additive in Kenya (Kipkore et al., 2014), and are regularly consumed with food in South Africa, Botswana and Zimbabwe (Bhebhe et al., 2016; Motlhanka and Makhabu, 2011; Shikanga et al., 2010). *L. javanica* leaves have ethnomedicinal repute for various conditions, including respiratory and gastrointestinal ailments, wound healing, and dysentery (de Wet et al., 2010; Kipkore et al., 2014; Maroyi, 2017; Pascual et al., 2001), and also has reported anti-parasitic and insecticidal uses (Lukwa, 1994; Madzimure et al., 2011). It recently gained popularity as tisane, prepared from a tea cut comprising dried stalk and leaves. International regulatory requirements for safe foodstuffs and topical applications demand risk assessment in various health endpoints to establish a safe toxicological profile from acute, subchronic and chronic use.

The composition of *L. javanica* has been studied by many (Asowata-Ayodele et al., 2016; Chagonda et al., 2000; Chagonda and Chalchat, 2015; Dlamini, 2006; Ludere et al., 2013; Lukwa et al., 2009; Narzary et al., 2015; Narzary and Basumatary, 2019; Olivier et al., 2010; Spittler, 2019; Viljoen et al., 2005). From its chemical characterisation, the tea cut composition raises no concerns from its phenolic, alkaloid, amino acid, mineral, metal and essential oil fractions, besides the presence of triterpenoid icterogenin, a reported hepatotoxin in sheep (Arias, 1963).

In recent years, cases of herb-induced liver injury from herbal food supplements/medicines have significantly increased, which in turn, elicit concern about consumption of herbal preparations. Considering the presence of icterogenin in *L. javanica* leaves, the dominating purpose of this study was to screen for the potential of hepatoxicity from consumption of *L. javanica* tisane.

In line with the 3 Rs of risk assessment (replacement, refinement and reduction of animals), rapid *in vitro* screening models offer paradigms for early prediction of hepatoxicity potential in herbal preparations. We used high content imaging technology with powerful image analysis software, which enables quantitative monitoring of morphological features in cultured cells indicative of different mechanisms of hepatotoxicity such as steatosis (Donato et al., 2012), oxidative stress, mitotoxicity and lysotoxicity (O’Brien et al., 2006; Persson et al., 2014; Tolosa et al., 2015; Zanella et al., 2010).

## 2. Methods

### 2.1. Materials and reagents

HepG2/C3A cells were purchased from Cellonex, South Africa; EMEM, FBS and MEM Non-essential amino acids (NEAA) from GE Healthcare Life Sciences (Logan, UT, USA); PBS with and without Ca2+ and Mg2+ and trypsin from Lonza (Walkersville, MD, USA); Bis-benzamide H 33342 trihydrochloride (Hoechst) and propidium iodide (PI) from Sigma (St. Louis, MO, USA). MitoTracker Green, LysoTracker Red, CellRox Orange, tetramethylrhodamine ethyl ester (TMRE), LipidTox Red for Neutral lipids, LipidTox Green phospholipidosis detection reagent and NucRed Live 647 were purchased from Molecular Probes – Life Technologies – Thermo Fisher Scientific (Logan, UT, USA).

### 2.2. Plant collection and extract preparation

Dried plant material of *L. javanica* (Burm. f.) Spreng (tea cut, code LJ-ZM-181218) was supplied by Parceval (Pty) Ltd (Wellington, South Africa). This material was wild harvested in Zimbabwe in December 2018. The name of the plant was verified using http://www.theplantlist.org which stated that the record derives from the WCSP where data was supplied on the 26 March 2012 and that the original publication details are as follows: *Syst. Veg. 2: 752 1825*. A tea extract of *L. javanica* was prepared by steeping 10 g of plant material in 1 L of boiled dH2O for 5 minutes. The extract was filtered using Whatman No. 1 filter paper, freeze dried and stored at 4°C in a desiccator until further use. A yield of 24.8% (g/100g of dry plant material) was obtained.

### 2.3. Cell seeding and treatment

C3A cells were maintained at 37°C in a humidified incubator with 5% CO2 in 10 cm culture dishes. Complete growth medium consisted of EMEM supplemented with 10% foetal bovine serum (FBS) (HyClone) and MEM Non-essential Amino Acids at 1X the stock concentration (HyClone).

C3A cells were seeded at 5000 cells/well of a 96 well plate in 100 µl aliquots and incubated at 37°C overnight. Treatment was done by adding an additional 100 µl of extract to achieve final concentrations of 12.5, 25, 50, 100 and 200 µg/ml, respectively. Cells were incubated at 37°C for 48 hours. Melphalan, at 0.781, 1.56, 3.13, 6.25 and 12.5 µM served as the positive control for cytotoxicity, reactive oxygen species production as well as mitochondrial content and membrane potential assays. Carbonyl cyanide m-chlorophenyl hydrazine (CCCP) at 1.56, 3.13, 6.25, 12.5 and 25 µM was included as secondary positive control for mitochondrial content and membrane potential assay. Chloroquine at 3.13, 6.25, 12.5, 25 and 50 µM served as the positive control for lysosomal content and neutral lipid detection assays.

### 2.4. Staining protocols

*All staining protocols:* Treatment medium was removed after 48 h and cells were washed with 100 µl of PBS containing Ca2+ and Mg2+. Hoechst 33342 diluted in DPBS to a final concentration of 5 µg/ml was added to wells using 100 µl aliquots as nuclear counterstain and incubated at 37°C for 30 minutes. The following additional stains were used to monitor cell death, ROS levels, mitochondrial health, neutral lipid content and lysosomal content: *Cytotoxicity:* PI was diluted in PBS to a concentration of 110 µg/ml and added to wells just before image acquisition using 10 µl aliquots. *ROS production:* CellRox Orange was diluted in 10 ml DPBS to a final concentration of 2.5 µM and 100 µl added to each well. Cells were incubated at 37°C for 30 minutes prior to imaging. *Mitochondrial content and membrane potential:* TMRE and MitoTracker Green were diluted together in DPBS to final concentrations of 0.25 µM and 0.02 µM. Cells were stained using 100 µl aliquots and incubated at 37°C for 30 minutes prior to imaging. *Neutral lipids:* Cells were fixed using 4% formaldehyde in DPBS for 15 minutes. Fixative was removed and cells washed twice with 100 µl aliquots of DPBS. The 1000X LipidTox Red neutral lipid stain was diluted 1:1000 in DPBS and added to wells using 100 µl aliquots. Cells were incubated at 37°C for 30 minutes prior to imaging. *Lysosomal content:* LysoTracker Red was diluted in DPBS to a final concentration of 0.05 µM and added to wells using 100 µl aliquots. Cells were incubated at 37°C for 30 minutes prior to imaging.

### 2.5. Image acquisition and analysis

All images were acquired using the ImageXpress Micro XLS Widefield High-Content Analysis System (Molecular Devices) with plate acquisition setup for a 96 well plate at a magnification of 10x and spatial distribution set at 9 sites per well (3 by 3). Combinations of DAPI, FITC, TRITC or Texas Red filters were used for hepatotoxicity assays depending on the dyes used in each assay and the Cy5 filter for micronucleus assay. Image analysis was performed using the Multi Wavelength Cell Scoring analysis module (hepatotoxicity) and Micronucleus module (genotoxicity) of MetaXpress version 6.1 High-Content Image Acquisition and Analysis Software. Analysis was based on segmentation parameters defined for each wavelength such as minimum and maximum widths, the intensity above local background and the minimum stained area of the cellular component of interest.

### 2.6. Statistical analysis

All experiments were repeated three times. The two-tailed Student t-test for two samples assuming equal variance was used to determine statistical significance. Error bars indicate the standard deviation of the mean for three experiments.

## 3. Results

All hepatotoxicity results of *L. javanica* as determined in the concentration range 12.5 – 200 µg/ml are summarized in Table 1. Indications of hepatotoxicity would include one or more of the following morphological changes, as observed for the respective positive controls (Table 2): increased ROS (oxidative stress), decreased mitochondrial membrane potential and mitochondrial content (mitotoxicity), increased lipid accumulation (steatosis), increased lysosomal content and/or increased cell death. The cytotoxicity results for *L. javanica* revealed no change in number of live (stained with Hoechst only) or dead cells (stained with Hoechst and PI). No significant changes were observed in ROS production, and no increase was observed in lysosomal content.

**Table 1:** Summary of the effects of *L. javanica* on morphological features in C3A cells.

| Parameter | *L. javanica* extract concentration (µg/ml) |  |  |  |  |  |
| --- | --- | --- | --- | --- | --- | --- |
|  | Control | 12.5 | 25 | 50 | 100 | 200 |
| Cytotoxicity (Number of live cells) | 2266 ± 159 | 2068 ± 308 | 2239 ± 259 | 2087 ± 121 | 2061 ± 308 | 1937 ± 300 |
| ROS production | 100.00 ± 0.68 | 98.18 ± 4.57 | 97.44 ± 3.03 | 97.01 ± 2.97 | 96.23 ± 3.06 | 98.05 ± 3.44 |
| Mitochondrial membrane potential | 100.00 ± 0.74 | 90.57 ± 6.01 | 88.53 ± 5.01 * | 90.99 ± 4.25 * | 90.49 ± 5.51 * | 91.01 ± 3.36 * |
| Mitochondrial content | 100.00 ± 8.29 | 105.84 ± 9.05 | 83.99 ± 4.93 | 94.68 ± 5.99 * | 86.41 ± 6.21 | 135.46 ± 15.36 * |
| Neutral lipid content | 100.00 ± 0.91 | 120.72 ± 14.82 | 87.35 ± 6.15 * | 80.26 ± 9.46 * | 77.56 ± 11.78 * | 70.70 ± 13.55 * |
| Lysosomal content | 100.00 ± 1.35 | 77.26 ± 4.57 # | 94.41 ± 2.87 * | 99.08 ± 3.77 | 99.36 ± 5.77 | 95.14 ± 5.99 |

\*p<0.05; #p<0.001 compared to control cells (blue font: reduced; red font: increased)

**Table 2:** Summary of the effects of positive controls on morphological features in C3A cells.

| Parameter | Melphalan (μM) |  |  |  |  |  |
| --- | --- | --- | --- | --- | --- | --- |
|  | Control | 0.78125 | 1.5625 | 3.125 | 6.25 | 12.5 |
| Cytotoxicity (Number of live cells) | 2244 ± 65 | 1532 ± 155 # | 1230 ± 167 # | 936 ± 94 # | 794 ± 72 # | 662 ± 45 # |
| ROS production | 100.00 ± 0.68 | 102.39 ± 2.79 | 104.15 ± 2.72 * | 107.51 ± 3.00 * | 112.58 ± 2.31 # | 121.75 ± 3.91 # |
| Mitochondrial membrane potential | 100.00 ± 0.74 | 98.62 ± 3.25 | 101.19 ± 2.11 | 111.30 ± 2.69 # | 118.76 ± 2.54 # | 123.70 ± 2.74 # |
| Mitochondrial content | 100.00 ± 8.29 | 203.55 ± 27.23 # | 232.97 ± 38.37 # | 318.56 ± 26.35 # | 389.14 ± 34.87 # | 449.62 ± 59.24 # |
| Parameter | CCCP (μM) |  |  |  |  |  |
|  | Control | 1.5625 | 3.125 | 6.25 | 12.5 | 25 |
| Mitochondrial membrane potential | 100.00 ± 0.74 | 92.67 ± 2.23 # | 82.16 ± 1.78 # | 66.18 ± 0.22 # | 60.42 ± 5.27 # | 53.25 ± 3.08 # |
| Mitochondrial content | 100.00 ± 8.29 | 85.64 ± 5.14 * | 78.47 ± 5.01 * | 63.46 ± 3.92 # | 70.90 ± 8.40 * | 77.09 ± 6.53 * |
| Parameter | Chloroquine (μM) |  |  |  |  |  |
|  | Control | 3.125 | 6.25 | 12.5 | 25 | 50 |
| Neutral lipid content | 100.00 ± 0.91 | 114.02 ± 10.46 | 107.34 ± 10.86 | 213.84 ± 24.74 * | 287.86 ± 25.63 # | 366.22 ± 30.58 # |
| Lysosomal content | 100.00 ± 1.35 | 293.30 ± 80.19 * | 412.68 ± 85.25 * | 1254.7 ± 141.72 # | 1406.4 ± 100.60 # | 1331.6 ± 130.85 # |

\*p<0.05; #p<0.001 compared to control cells (blue font: reduced; red font: increased)

The only parameter that revealed any potential hepatotoxic response from this extract was the mitochondrial membrane potential, where all concentrations from 25 µg/ml and higher reduced the membrane potential by around 10% compared to the control (p<0.05).

## 4. Discussion

No human, but limited animal studies were found relating *L. javanica* to toxic effects from its exposure. Adverse neurological effects were shown in mice from acute administration of high doses (4 ml of 12.55 - 37.5% (v/v)) of aqueous leaf extracts. Lethargy, observed in all animals and at all doses, were attributed to xanthine, (present at low levels in leaves), which were argued to produce vasodilation and overstimulation, leading to depression (Madzimure et al., 2011). Haematological and biochemical analyses were not performed. In another animal study, high doses of aqueous leaf extracts (450 - 1000 mg/kg bw/d for 28 days) were administered orally and intraperitoneally to mice for observation of adverse effects in various health endpoints, including haematological and biochemical parameters being relevant to this study. Statistically significant dose-dependent changes were observed in total bilirubin levels (decrease) and in percent liver to body weight relative to the control (increase). The observed hepatocellular injury was attributed to cytotoxic effects from exposure to high doses of extract constituents. Whilst the study did not attempt to present a causal effect, given the presence of icterogenin in the leaves, the findings are not unexpected for high dose exposure. However, consumption of *L. javanica* tisane implies low-rate exposure to potential hepatoxins.

The use of *L. javanica* leaves in a broiler diet was evaluated as a growth stimulation alternative to prophylactic antibiotics(Mpofu et al., 2016). Satisfactory growth, without reported adverse effects were observed over the 45-day diet at doses of 5 and 12g/kg. However, at the time, toxicity investigations were not a focus. In a follow-up study, *L. javanica*, fed to broiler chickens (63-days at 25 and 50 g/kg), evaluated blood chemistry parameters and reported significant changes were observed in bilirubin, alanine aminotransferase (ALT), aspartate transaminase (AST), sodium, potassium, cholesterol and magnesium levels (Mnisi et al., 2017). In another 9-week high-dose feeding study of Japanese quails no significant impact were observed in carcass characteristics and internal organs, but increases were observed in gizzard weights and ALT levels; Significant increase in meat yellowness was also observed post slaughter (Mnisi et al., 2017).

The highest concentration used in our investigation was based on the consumption of 2 cups (500 ml) of *L. javanica* tea (1.24 g dried extract) per day, and assuming 100% bioavailability as worst-case-scenario. Given a typical blood volume of 6 L, implies 207 µg/ml *L. javanica* in the blood for the given parameters. Results showed an absence of any dose-dependent changes in parameters that represent oxidative stress, steatosis, or lysosomal toxicity. Steatosis can be induced through interference with several metabolic pathways such as fatty acid beta-oxidation in mitochondria, fatty acid biosynthesis and others. Characterized by an increase in neutral lipid accumulation in hepatocytes, steatosis could lead to liver failure. The lack of dose-dependent responses suggests a low risk of steatosis.

The observed reduction (∼10%) in mitochondrial membrane potential was accompanied by a significant increase in mitochondrial content at 200 µg/ml, with an insignificant reduction in live cell numbers (p>0.05). Given the strong evidence of dose-dependent effects on the causal relationship between exposure and outcome, the observed result at this specific mid-range concentration is not considered as an important indicator of potential liver injury at this dose.

The dose-dependent decrease observed in neutral lipid content in response to treatment with *L. javanica* from 25 µg/ml and higher (Table 1) might be indicative of potential hepatoprotective effects at this intake level. Others already alluded to hepatoprotective effects from normal *L. Javanica* intake (Maroyi, 2017; Tapas et al., 2008).

## 5. Conclusion

Although the HepG2/C3A hepatocarcinoma cell line is frequently used for predictive hepatotoxicity screening, *in vitro* studies using cancer cells should be interpreted with caution. Rapid screening and evaluation of herb-induced liver injury from consumption of *L. javanica* leaf extracts suggest that the material is not anticipated to be hepatotoxic at low dose, i.e. typically 1-2 cups of tea per day. Screening tests present a first step in continued development of drugs or other preparations. However, considering of the observed effects at high doses, more detailed studies would be required for higher intake doses of this material, Finally, in view of the findings from this study and some of the reported ethnomedicinal uses, the results may also be extended to topical applications, hence applied to foods, medicines, and cosmetics. Given that topical application implies both reduced systemic availability and avoidance of first-pass metabolism of the extract, results suggest the risk of hepatoxicity from topical use of *L. javanica* leaf extracts is likely insignificant.

## Author contributions

M.v.d.V., T.C.K., L.V. and E.C. conceived and designed the experiments; B.S. performed the experiments; B.S. and M.v.d.V. analyzed the data; E.C., B.S., N.P.K. and M.v.d.V. wrote the paper; All authors have read and approved the manuscript.

## Appendix A. Supplementary data

LC-MS analysis data of the extract is provided in the Supplementary data file.

