## Supplemental data for "*In vitro* hepatotoxicity assessment of *Lippia javanica* (Burm.f.) Spreng. aqueous leaf extract"

**Supplementary data: LC-MS analysis on *Lippia javanica***

LC-MS analysis was performed at the Technische Universität Dortmund, Germany and the methods and results below were copied from their report.

**Sample preparation**

About 5.0 g powder of the dried sample (Table S1) was extracted with 50 mL methanol for 24 hr and sonicated at room temperature for 15 min. The extract was filtered and filled to 50 ml with methanol prior to analysis. LCMS analysis was done using the Luna C18 (3 µm) 50x3 mm column with injection volume 5 µL.

**Table S1: Sample information**

| Harvest date | Qty | Origin | Plant Part | Code | Notes |
|---|---|---|---|---|---|
| Dec-18 | 2 x 100g | Zimbabwe | Leaf | LJ-ZM-181218 | Wild harvested |

**LCMS analytical method**

High-resolution mass spectra (ESI-HRMS) were carried out on a LTQ Orbitrap spectrometer (Thermo Scientific, USA) equipped with a HESI-II source. The spectrometer was equipped with an Agilent 1200 HPLC system (Santa Clara, USA) including pump, PDA detector, column oven (30°C) and auto-sampler (injection volume: 5 µL for Fullscan, 7µL for $MS^n$). The HPLC analyses were performed with a Luna C18 column (50 x 3 mm, 3 µm particle size) from Phenomenex (Torrance, USA) with water (+ 0.1% formic acid) (A) and methanol (B) gradient (flow rate 350 µL/min). The gradient was set as follows: linear gradient from 95% A to 80% B over 49 min, 100% B isocratic for 5 min, the system returned within 0.5 min to initial conditions of 95% A and was equilibrated for 4.5 min.

**Results**

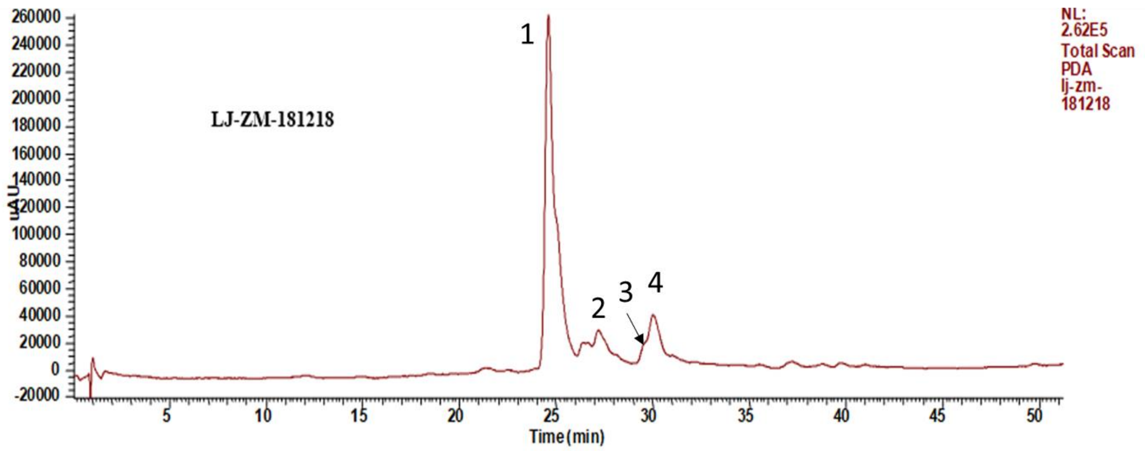


Figure S1: LCMS chromatogram of LJ-ZM-181218

**Table S2: Identification of the chemical constituents of *Lippia javanica* extract by LC–ESI–MS$^{n}$ analysis (refer to Appendices)**

| Peak no. | Rt (min) | Formula [M+H]$^{+}$ | Positive ion (*m/z*) | Ms2 | Proposed structure |
|---|---|---|---|---|---|
| 1 | 24.5 | $C_{21}H_{27}O_{12}$ | 471.1494 | 325.09, 307.08, 163.03 | 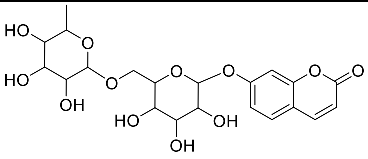 |
| 2 | 27.1 | $C_{21}H_{19}O_{12}$ | 463.0865 | 287.05 | 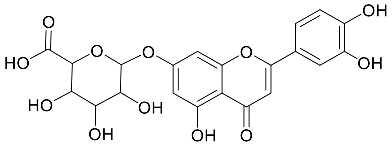 Luteolin 7-O-glucuronide |
| 3 | 30.0 | $C_{22}H_{21}O_{12}$ | 477.1020 | 301.07, 286.04, 258.05 | 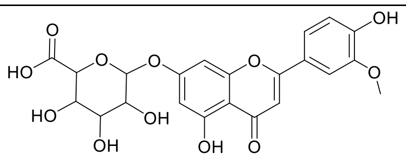 |
| 4 | 30.5 | $C_{23}H_{23}O_{13}$ | 507.1127 | 331.08 | 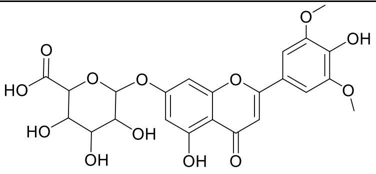 Tricin 7-O-glucuronide |

**Quantification**

**Hot water Extract**: About 2.0 g powder of each dried sample of *Lippia javanica* was soaked in 200 mL hot water (>80 °C) for 5 min. The extracts were filtered and lyophilized. The extract was dissolved in 40 ml methanol prior to analysis.

For LCMS analysis, 100 μL of the extract mixed with 10 μL of internal standard $^{13}C_3$-Catechin (50 μg/mL, $C_{12}{}^{13}C_3H_{15}O_6$). The analyses were done using the Luna C18 (3 μm) 50x3 mm column with injection volume was 5 μL.

Compounds 1-4 were quantified with available reference compound rutin (Table S3).

**Table S3: Content of compounds 1-4 (μg/g d.w.)**

| Sample code | **1**<br>471.1494<br>$C_{21}H_{27}O_{12}$ | **2**<br>463.0865<br>$C_{21}H_{19}O_{12}$ | **3**<br>477.1020<br>$C_{22}H_{21}O_{12}$ | **4**<br>507.1127<br>$C_{23}H_{23}O_{13}$ |
|---|---|---|---|---|
| LJ-ZM-181218 | 2373.8 | 1960.1 | 2416.0 | 541.1 |

### Appendices

#### Compound **1**

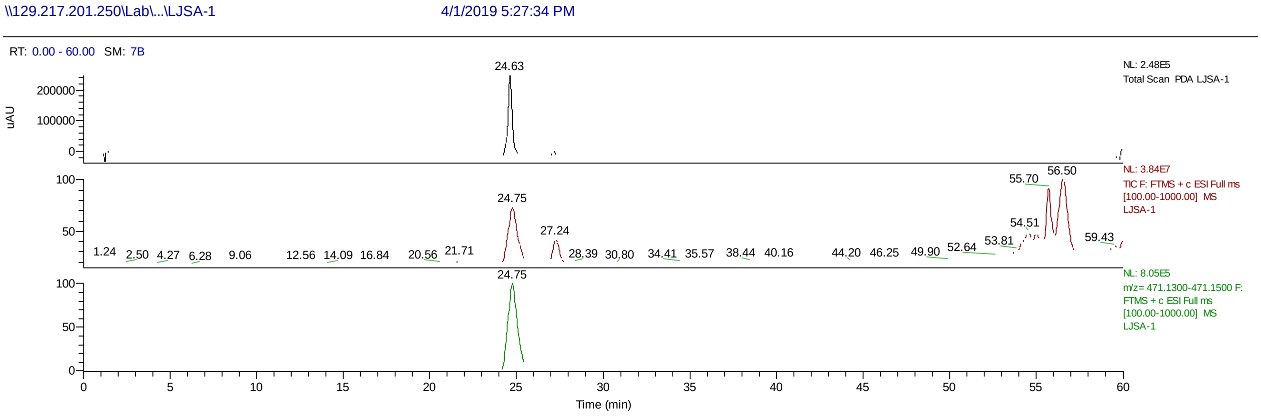


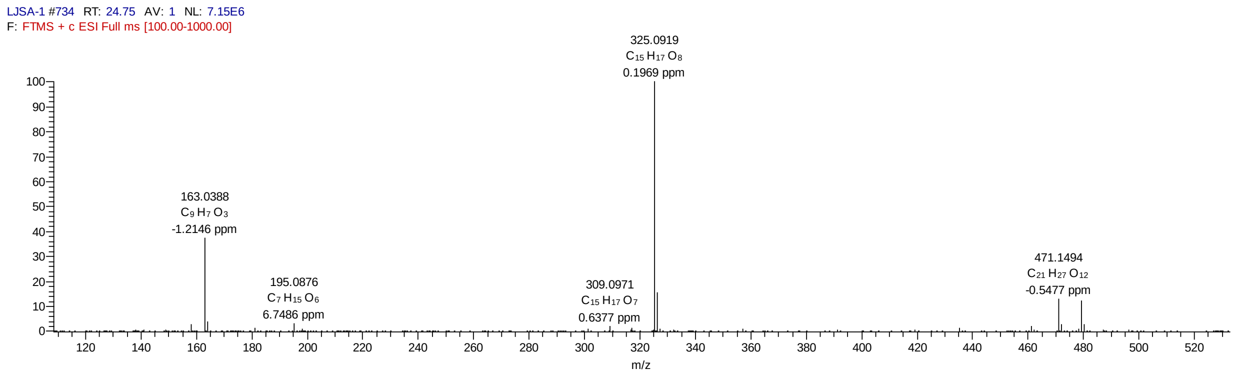


ms2@325.09

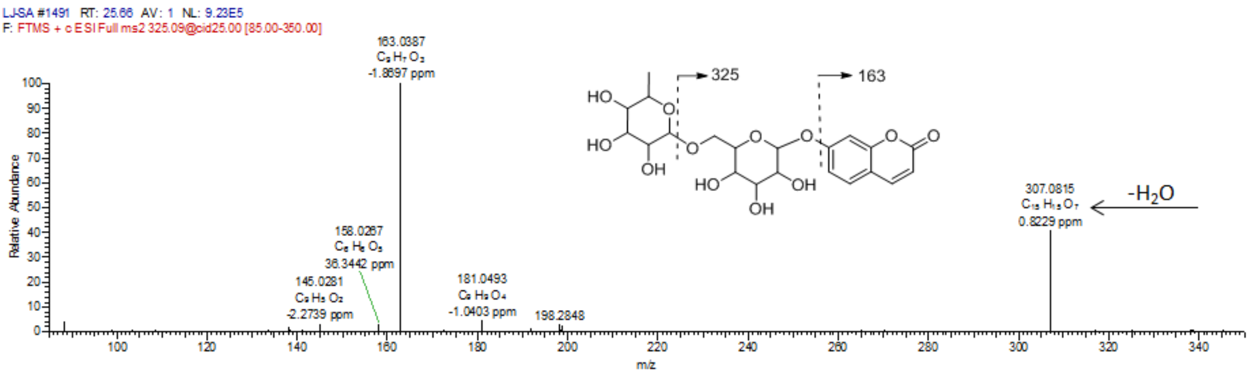

### Compound **2**

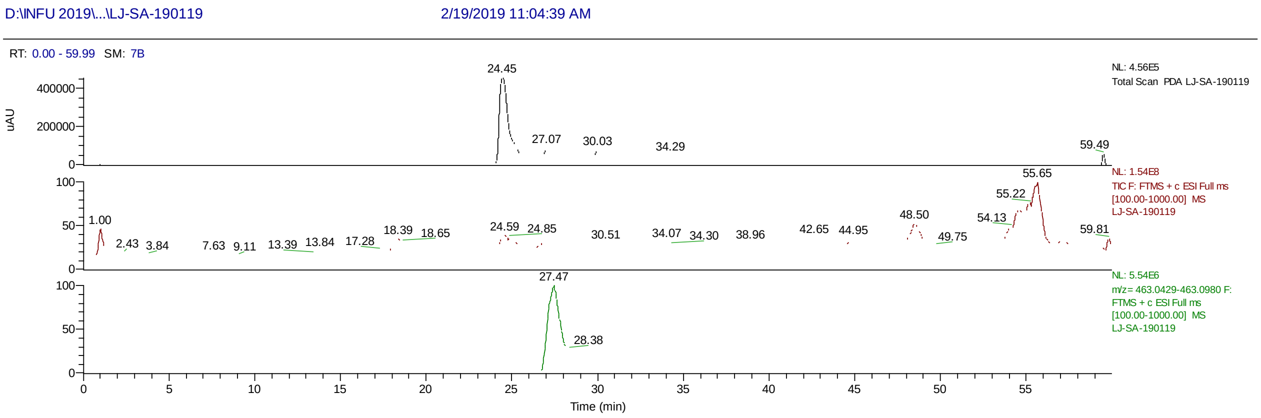

D:\INFU 2019\...\LJ-SA-190119
2/19/2019 11:04:39 AM
RT: 0.00 - 59.99 SM: 7B
NL: 4.56E5
Total Scan PDA LJ-SA-190119
NL: 1.54E8
TIC F: FTMS + c ESI Full ms [100.00-1000.00] MS LJ-SA-190119
NL: 5.54E6
m/z= 463.0429-463.0980 F: FTMS + c ESI Full ms [100.00-1000.00] MS LJ-SA-190119
uAU
Time (min)


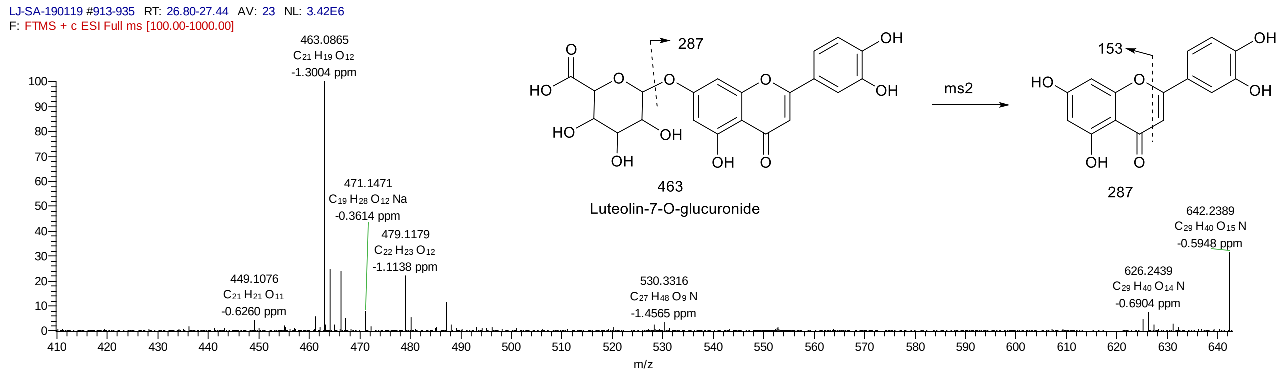

LJ-SA-190119 #913-935 RT: 26.80-27.44 AV: 23 NL: 3.42E6
F: FTMS + c ESI Full ms [100.00-1000.00]
463.0865
C21 H19 O12
-1.3004 ppm
471.1471
C19 H28 O12 Na
-0.3614 ppm
479.1179
C22 H23 O12
-1.1138 ppm
449.1076
C21 H21 O11
-0.6260 ppm
530.3316
C27 H48 O9 N
-1.4565 ppm
626.2439
C29 H40 O14 N
-0.6904 ppm
642.2389
C29 H40 O15 N
-0.5948 ppm
287
463
Luteolin-7-O-glucuronide
ms2
153
287
m/z


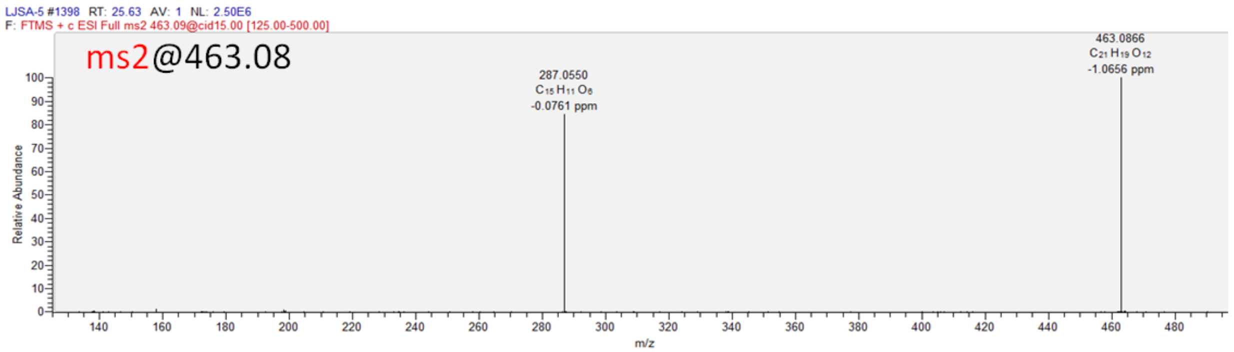

LJSA-5 #1398 RT: 25.63 AV: 1 NL: 2.50E6
F: FTMS + c ESI Full ms2 463.09@cid15.00 [125.00-500.00]
ms2@463.08
287.0550
C15 H11 O6
-0.0761 ppm
463.0866
C21 H19 O12
-1.0656 ppm
Relative Abundance
m/z


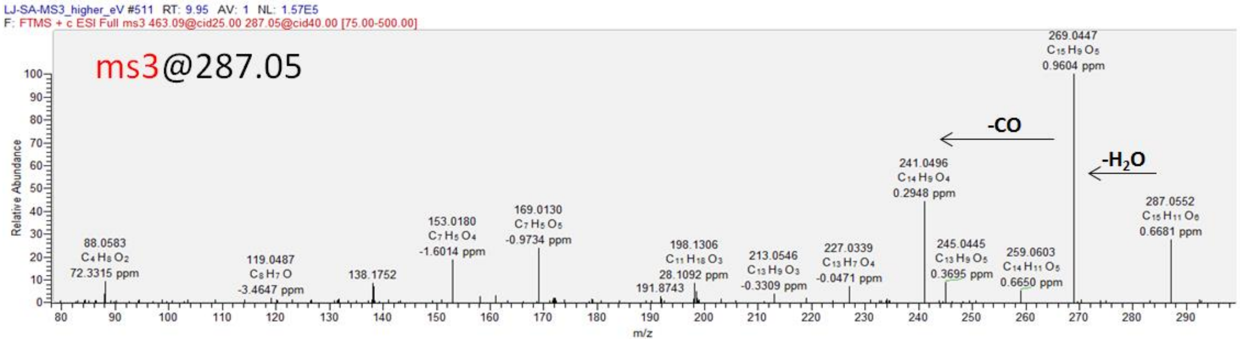

LJ-SA-MS3_higher_eV #511 RT: 9.95 AV: 1 NL: 1.57E5
F: FTMS + c ESI Full ms3 463.09@cid25.00 287.05@cid40.00 [75.00-500.00]
ms3@287.05
269.0447
C15 H9 O5
0.9604 ppm
-CO
-H2O
241.0496
C14 H9 O4
0.2948 ppm
287.0552
C15 H11 O6
0.6681 ppm
88.0583
C4 H8 O2
72.3315 ppm
119.0487
C8 H7 O
-3.4647 ppm
138.1752
153.0180
C7 H5 O4
-1.6014 ppm
169.0130
C7 H5 O5
-0.9734 ppm
198.1306
C11 H18 O3
28.1092 ppm
191.8743
213.0546
C13 H9 O3
-0.3309 ppm
227.0339
C13 H7 O4
-0.0471 ppm
245.0445
C13 H9 O5
0.3695 ppm
259.0603
C14 H11 O5
0.6650 ppm
Relative Abundance
m/z

Compound **3**

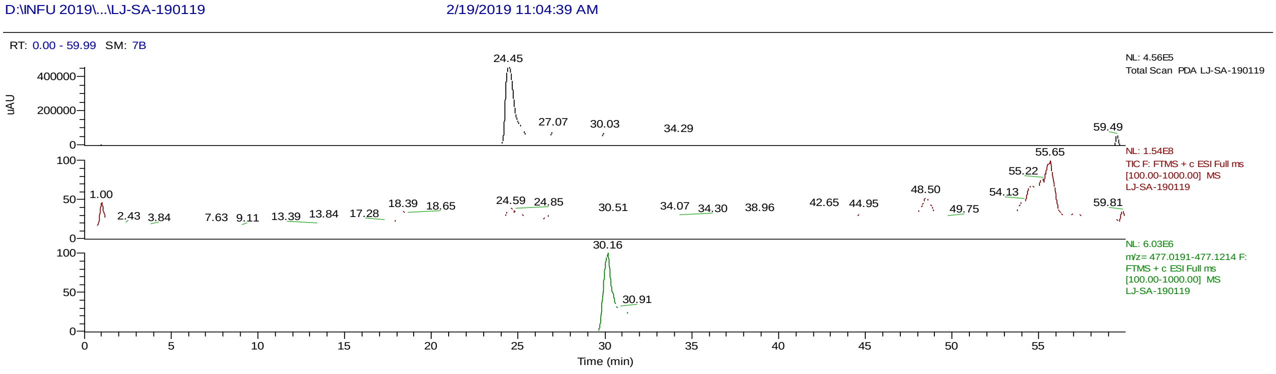

D:\INFU 2019\...\LJ-SA-190119
2/19/2019 11:04:39 AM
RT: 0.00 - 59.99 SM: 7B
NL: 4.56E5
Total Scan PDA LJ-SA-190119
NL: 1.54E8
TIC F: FTMS + c ESI Full ms [100.00-1000.00] MS LJ-SA-190119
NL: 6.03E6
m/z= 477.0191-477.1214 F: FTMS + c ESI Full ms [100.00-1000.00] MS LJ-SA-190119
uAU
Time (min)


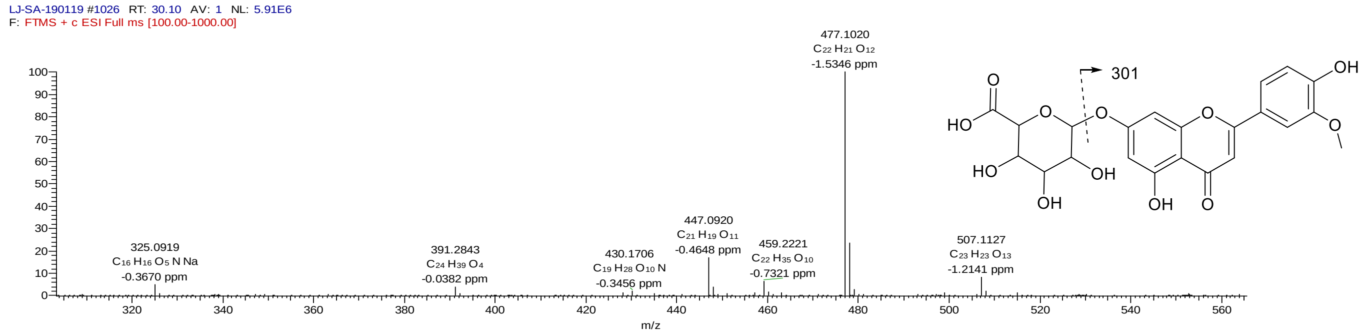

LJ-SA-190119 #1026 RT: 30.10 AV: 1 NL: 5.91E6
F: FTMS + c ESI Full ms [100.00-1000.00]
477.1020
C22 H21 O12
-1.5346 ppm
325.0919
C16 H16 O5 N Na
-0.3670 ppm
391.2843
C24 H39 O4
-0.0382 ppm
430.1706
C19 H28 O10 N
-0.3456 ppm
447.0920
C21 H19 O11
-0.4648 ppm
459.2221
C22 H35 O10
-0.7321 ppm
507.1127
C23 H23 O13
-1.2141 ppm
m/z
301


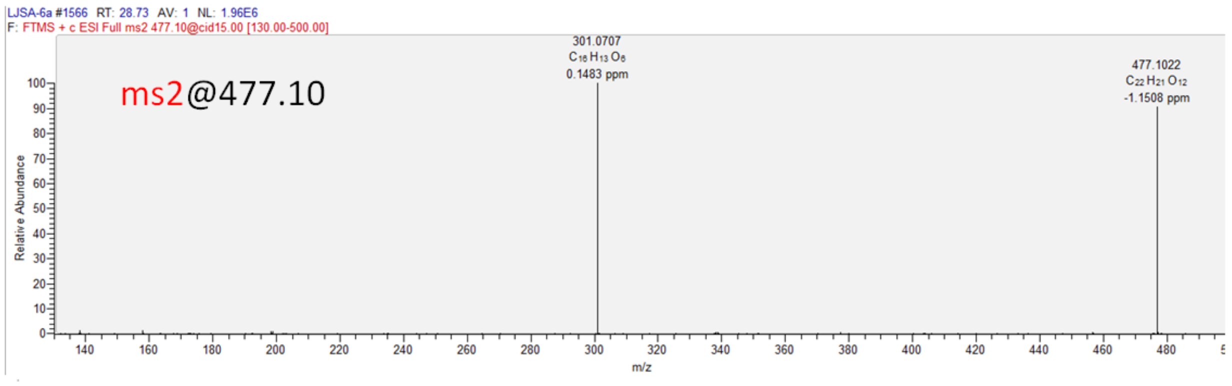

LJSA-6a #1566 RT: 28.73 AV: 1 NL: 1.96E6
F: FTMS + c ESI Full ms2 477.10@cid15.00 [130.00-500.00]
ms2@477.10
301.0707
C16 H13 O6
0.1483 ppm
477.1022
C22 H21 O12
-1.1508 ppm
Relative Abundance
m/z


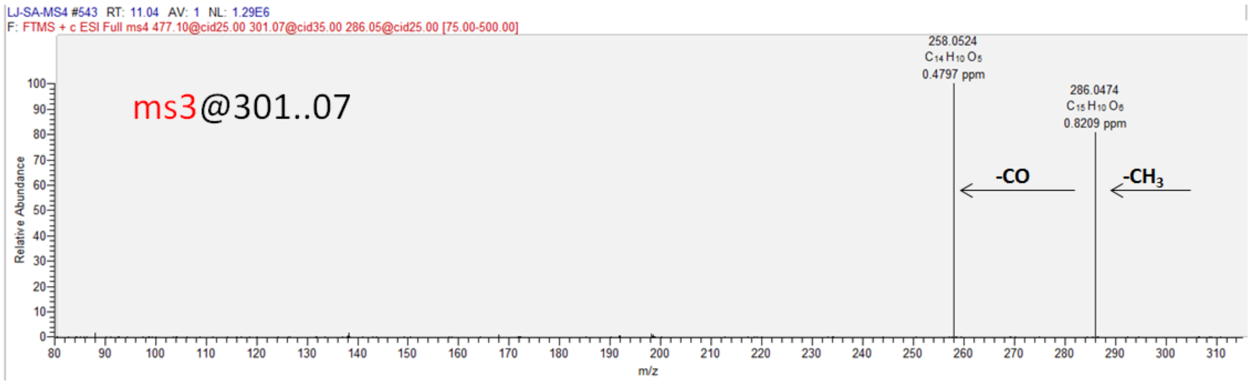

LJ-SA-MS4 #543 RT: 11.04 AV: 1 NL: 1.29E6
F: FTMS + c ESI Full ms4 477.10@cid25.00 301.07@cid35.00 286.05@cid25.00 [75.00-500.00]
ms3@301..07
258.0524
C14 H10 O5
0.4797 ppm
286.0474
C15 H10 O6
0.8209 ppm
-CO
-CH3
Relative Abundance
m/z

### Compound **4**

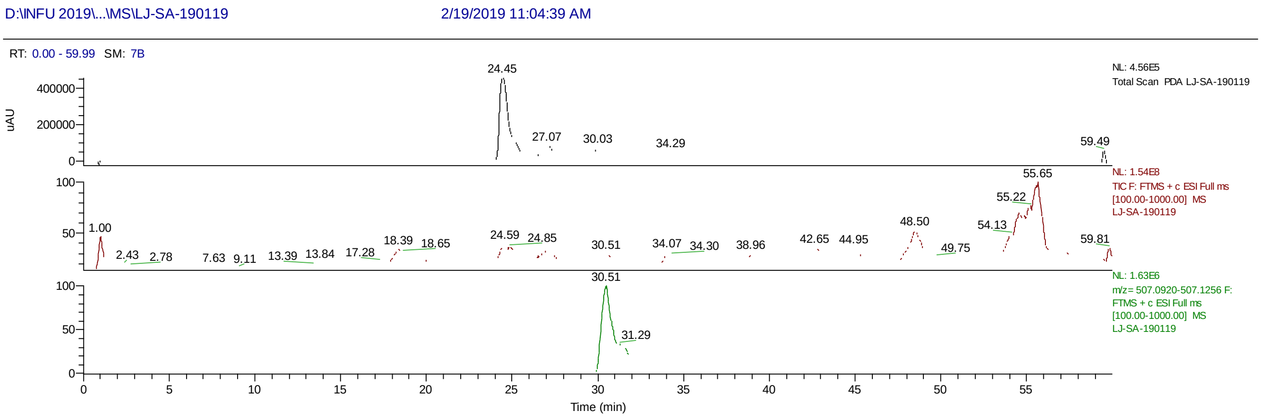


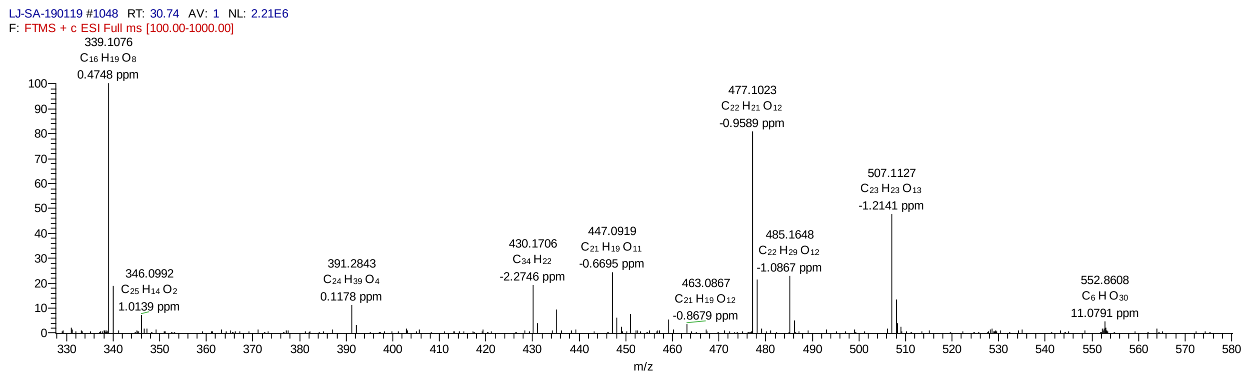


ms2@507.11

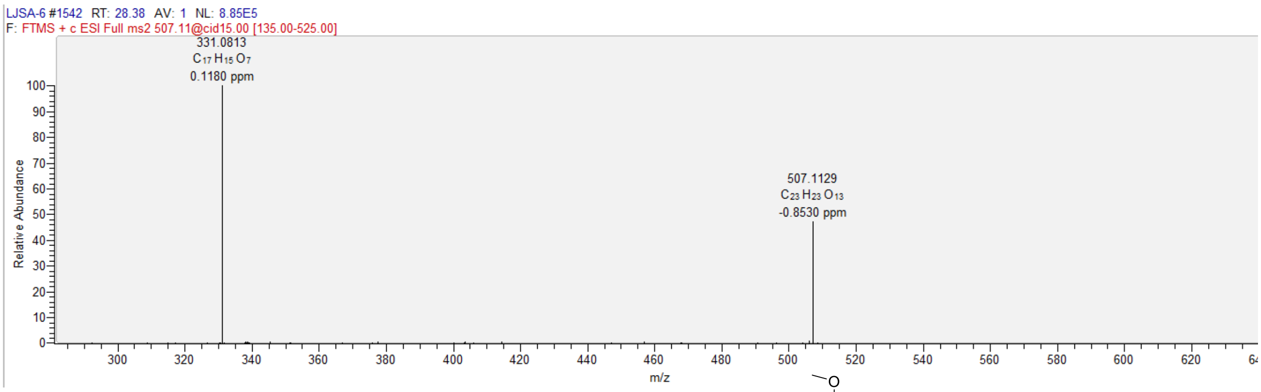


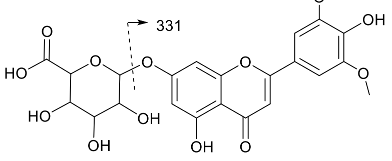


Tricin 7-O-glucuronide
